# Characterization of the *Corynebacterium glutamicum* dehydroshikimate dehydratase QsuB and its potential for microbial production of protocatechuic acid

**DOI:** 10.1101/2020.03.27.011478

**Authors:** Ekaterina A. Shmonova, Olga V. Voloshina, Maksim V. Ovsienko, Sergey V. Smirnov, Vera G. Doroshenko

**Author notes:** Corresponding author; (VGD).

## Abstract

The dehydroshikimate dehydratase (DSD) from *Corynebacterium glutamicum* encoded by the *qsuB* gene is related to the previously described QuiC1 protein (39.9% identity) from *Pseudomonas putida*. QuiC1 and QsuB are both two-domain bacterial DSDs. The N-terminal domain provides dehydratase activity, while the C-terminal domain has sequence identity with 4-hydroxyphenylpyruvate dioxygenase. Here, the QsuB protein and its DSD domain (N-QsuB) were expressed in the T7 system, purified and characterized. QsuB was present mainly in octameric form (60%), while N-QsuB had a predominantly monomeric structure (80%) in solution. Both proteins possessed DSD activity with one of the following cofactors (listed in order of decreasing activity): Co^2+^, Mg^2+^, Mn^2+^ or Ca^2+^. The K_*m*_ and k_*cat*_ values for QsuB were two and three times higher, respectively (K_*m*_ ~ 1 mM, k_*cat*_ ~ 61 s^−1^) than those for N-QsuB. Notably, 3,4-DHBA inhibited both enzymes via an uncompetitive mechanism. QsuB and N-QsuB were tested for 3,4-DHBA production from glucose in *E. coli*. MG1655Δ*aroE* P_*lac*_–*qsuB* produced at least two times more 3,4-DHBA than MG1655Δ*aroE* P_*lac*_–*n*-*qsuB* in the presence of isopropyl β-D-1-thiogalactopyranoside.

## Introduction

Protocatechuic acid or 3,4-dihydroxybenzoic acid (3,4-DHBA) is a catechol-type phenol present in some fruits and vegetables [1, 2]. It has been shown that 3,4-DHBA displays antibacterial [3], antimutagenic [4], anti-inflammatory [5], antihyperglycemic [6] and highly antioxidant [7] effects. It also has potential applications for further microbiological synthesis of numerous valuable compounds, including the bioplastic precursor *cis,cis*-muconic acid [8, 9]. To produce 3,4-DHBA from glucose, the synthesis of this compound from DHS, an intermediate in the common aromatic pathway, was previously implemented in *E. coli* cells [10, 11]. This reaction was catalyzed by dehydroshikimate dehydratase (DSD) (EC: 4.2.1.118) (Fig 1), a part of the quinate and shikimate degradation pathways in soil-dwelling bacteria and fungi. Another source of this enzyme is the biosynthesis of the unique 3,4-catecholate moiety found as part of the petrobactin scaffold present in the *Bacillus cereus* group. DSD is encoded by the *asbF* gene and, being identical in *Bacillus anthracis* and *Bacillus thuringiensis*, has been thoroughly investigated [12, 13]. Based on primary structure analysis, all known DSDs were divided into four groups: fungal single-domain, bacterial two-domain, AsbF-like, and bacterial membrane-associated enzymes [14]. The application of AsbF and fungal (*Podospora pauciseta*) and bacterial two-domain (*Klebsiella pneumoniae*) enzymes to obtain 3,4-DHBA as an intermediate for the generation of valuable compounds from glucose has been demonstrated [10, 11, 15, 16].

**Fig 1.**
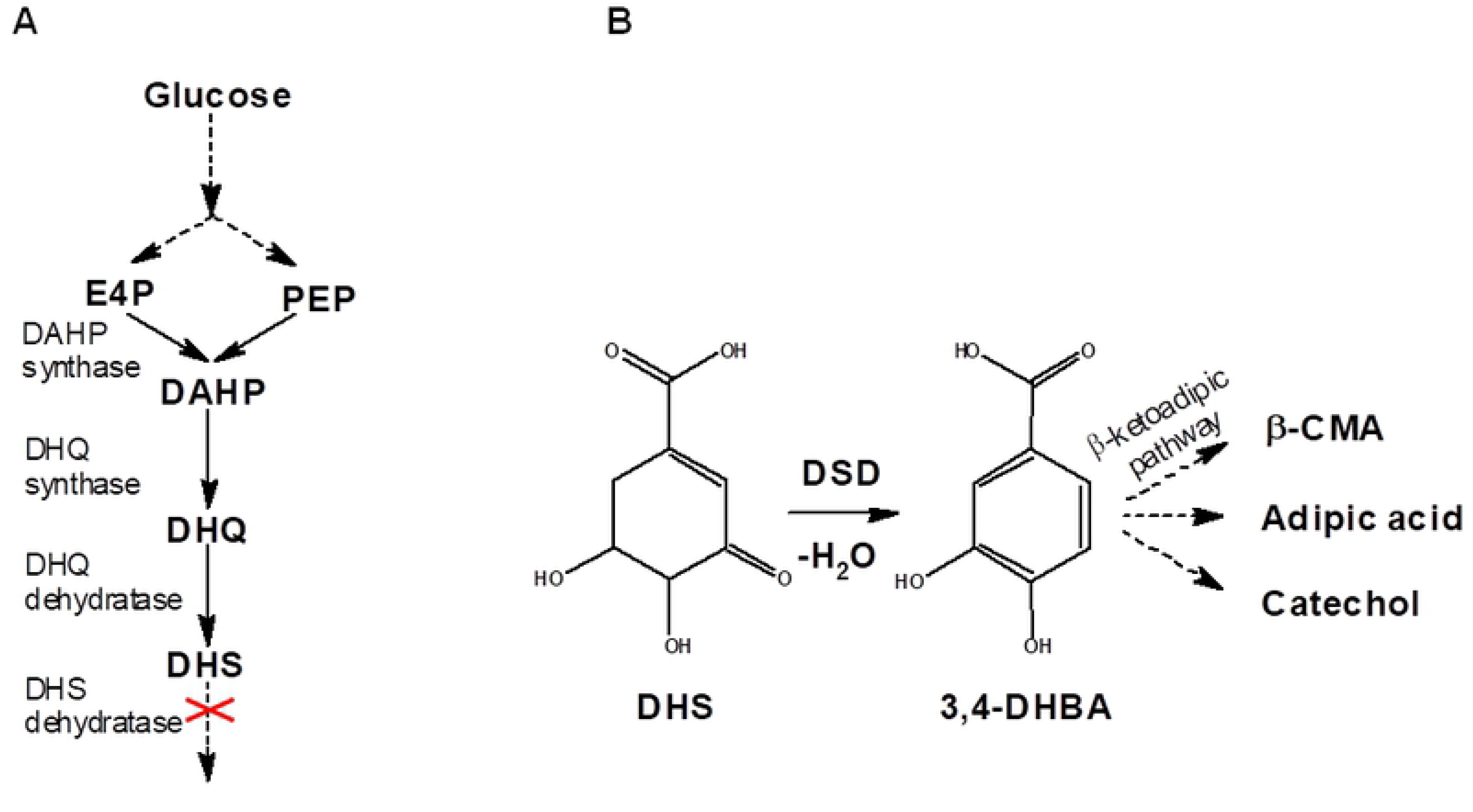
Biosynthesis of 3,4-DHBA from 3-dehydroshikimic acid (DHS), an intermediate of the common aromatic pathway. (A) Synthesis of DHS from glucose. (B) Formation of 3,4-DHBA from DHS with the help of DSD. Abbreviations: β-CMA - β-carboxy-*cis,cis*-muconic acid; DAHP - 3-deoxy-D-arabino-heptulosonate-7-phosphate; DHQ - 3-dehydroquinate; E4P - erythrose 4-phosphate; PEP - phosphoenolpyruvate.

The well-known industrial microorganism *Corynebacterium glutamicum*, which has generally recognized as safe (GRAS) status, harbors DSD, which has not yet been characterized biochemically. DSD is encoded by the *qsuB* gene, a part of the operon *qsuABCD*, whose products are involved in the quinate/shikimate utilization pathway [17]. Previously, QsuB of *C. glutamicum* was assigned to the two-domain bacterial DSD group [14]. QuiC1 from *Pseudomonas putida* was the only characterized member of this class. The N-terminal domain of QuiC1 catalyzes a dehydratase reaction, while the C-terminal domain has significant sequence identity with 4-hydroxyphenylpyruvate dioxygenase (HPPD) (EC: 1.13.11.27). The role of the C-terminal domain is still unknown, but it was suggested that it might be involved in 3,4-DHBA degradation via the ß-ketoadipate pathway. Here, we examined the biochemical characteristics of full-length QsuB and its N-terminal domain (N-QsuB) as well as QsuB application to 3,4-DHBA production in heterologous hosts.

## Materials and methods

### Bacterial strains and growth conditions

Known laboratory *E. coli* strains [18]: MG1655 (F^−^λ^−^*ilvG rfb*-50 *rph*-1) and BL21(DE3) (F^−^*ompT gal dcm lon hsdS_B_*(r_B_^−^m_B_^−^) λ(DE3[*lacI lacUV5-T7p07 ind1 sam7 nin5*]) [*malB^+^*]_K-12_(λ^S^) were subjected to further modifications in this work. The strains were cultivated in rich LB medium [18]. For the preparation of electrocompetent *E. coli* cells, SOB medium was used [19]. Antibiotics, when required, were added in the following concentrations (mg/l): ampicillin – 200, chloramphenicol – 20, tetracycline – 12.5.

Cell cultures for the isolation of the QsuB and N-QsuB proteins were prepared as follows. Flasks containing 30 ml LB with ampicillin were inoculated with overnight cultures (300 μl) of the BL21(DE3)/pET22b-*qsuB* and BL21(DE3)/pET22b-*n*-*qsuB* strains. The flasks were incubated at 25°C and 200 rpm for two h, then subjected to 1 mM isopropyl β-D-1-thiogalactopyranoside (IPTG) induction and incubated for an additional 20 h.

Fermentations were performed in tubes (18×200 mm) containing 2 ml of the production medium: 40 g/l glucose, 60 g/l CaCO_3_, 10 g/l tryptone, 10 g/l NaCl, 5 g/l yeast extract, 0.5 g/l (NH_4_)_2_SO_4_, 0.5 g/l K_2_HPO_4_, 5 mg/l FeSO_4_ × 7H_2_O, 4 mg/l MnSO_4_ × 5H_2_O, 10 mg/l thiamine, 10 mg/l 4-hydroxybenzoic acid, 10 mg/l 4-aminobenzoic acid, and 10 mg/l 2,3-dihydroxybenzoic acid. The fermentation tubes were inoculated with 0.2 ml of seed culture, which was prepared as follows. One loop (3 mm) of cells from a fresh plate was inoculated into a tube (13×150 mm) containing 3 ml of LB and incubated at 34°C with aeration (240 rpm) for 3 h. The fermentation tubes were cultivated at 34°C (240 rpm) for 44 h. Then, DHS and 3,4-DHBA concentrations were determined using HPLC.

### DNA manipulation

All recombinant DNA manipulation was conducted according to standard procedures [18] and the recommendations of the enzyme manufacturer (Thermo Scientific, USA). Plasmid and chromosomal DNAs were isolated with Plasmid Miniprep (Evrogen, Russia) and PurElute Bacterial Genomic Kit (Edge BioSystems, USA), respectively. PCR was performed with Taq DNA polymerase (GBM, Russia) and with Phusion DNA Polymerase (Thermo Scientific) for circular polymerase extension cloning (CPEC) [20]. Primers (S1 Table) were purchased from Evrogen. Plasmids and genetic modifications of the *E. coli* chromosome were verified by sequence analysis.

### Plasmid construction

The plasmids pET22b-*qsuB* and pET22b-*n*-*qsuB* were obtained by CPEC followed by selection in BL21(DE3) cells. The DNA fragment containing pET22b (Novagen, USA) was amplified using the primers P1/P2. The DNA fragments of *qsuB* and *n*-*qsuB* were amplified using the chromosomal DNA of *C. glutamicum* 2256 ATCC13869 as a template and the primers P3/P4 and P3/P5, respectively.

To integrate the *qsuB* gene into the *E. coli* chromosome, a DNA fragment containing P_*lacUV5*_-*qsuB* was cloned into the integrative vector pAH162-λ*attL*-*tet*-λ*attR* [21] between the *Sal*I and *Sac*I restriction sites. The DNA fragment containing P_*lacUV5*_-*qsuB* was obtained by overlapping PCR (primers P6/P9) of DNA fragments containing the promoter P_*lacUV5*_ and _the_ *qsuB* coding region. These DNA fragments were amplified by PCR with the primers P6/P7 and P8/P9 using pELAC [22] and pET22b-*qsuB* plasmid DNA as templates, respectively.

### Strain construction

To construct the MG1655Δ*aroE* strain, an inframe deletion of the *aroE* gene was created using λRed-mediated integration of a DNA fragment containing the excisable marker λ*attL*-*cat*-λ*attR* (primers P10, P11). The recombinant plasmids pKD46 and pMW118-λ*attL*-*cat*-λ*attR* were used as a helper for the λRed-mediated integration and as a template for the excisable marker λ*attL*-*cat*-λ*attR*, respectively [19, 23]. The marker was excised using the helper plasmid pMW-*int*-*xis* [23].

The MG1655Δ*aroE*P_*lac*_-*qsuB* strain was created via integration of P_*lacUV5*_-*qsuB* into the native φ80-*attB* site of the MG1655 chromosome. The integrative vector pAH162-λ*attL*-*tet*-λ*attR* with the cloned modification P_*lacUV5*_-*qsuB* (see above) containing the φ80-*attP*-site and the pAH123-helper plasmid were used [21].

The MG1655Δ*aroE*P_*lac*_-*n*-*qsuB* strain was generated via λRed integration of the excisable marker λ*attL*-*cat*-λ*attR* instead of the 5’-part of the *qsuB* coding region (primers P12/P13) into the chromosome of the MG1655Δ*aroE*P_*lac*_-*qsuB* strain, followed by marker removal.

### Obtaining crude extracts and purified proteins

All manipulations were performed at 4°C. After cultivation, cells were collected by centrifugation at 13200 rpm for 5 min and washed twice with 0.9% NaCl. The pellets were resuspended in 0.5 ml of 0.1 M potassium phosphate (pH 7.5), 0.1 mM EDTA, and 0.4 mM dithiothreitol buffer solution with the addition of 0.1 mM phenylmethylsulfonyl fluoride. The obtained suspensions were sonicated and then centrifuged at 13200 rpm for 5 min. The supernatants were decanted and then subjected to enzymatic reactions and 12% SDS-PAGE. PageRuler Prestained Protein Ladder 26616 (Thermo Scientific) was used to evaluate protein molecular mass.

For protein purification, the cells were resuspended in 15 ml of the abovementioned buffer and lysed with a French press. The cell debris was eliminated by centrifugation at 6000 rpm for 5 min. Then, the supernatant was applied to Ni-NTA Affinity Resin (Clontech, USA). The protein fractions were eluted with an imidazole gradient from 20 mM (in the binding buffer) to 500 mM (in the elution buffer). The fraction containing the purified protein was dissolved in a buffer with 50 mM Tris-HCl (pH 7.5), 500 mM NaCl and 5% glycerol.

### Gel filtration

Molecular mass determination was carried out using gel-filtration chromatography with a Superose 6 Increase 10/300 GL (GE Healthcare, USA) column. A buffer solution containing 50 mM sodium phosphate (pH 7.0), 150 mM NaCl and a flow rate of 0.5 ml/min was used for N-QsuB. A buffer solution containing 50 mM Tris-HCl (pH 7.5), 500 mM NaCl, and 5% v/v glycerol and a flow rate of 0.4 ml/min were used for QsuB. Proteins were detected by monitoring the absorbance at 280 nm. The gel filtration markers MWGF 1000 (29 - 700 kDa) and MWGF 70 (12.4 kDa) were purchased from Sigma-Aldrich, USA.

### Measurement of DSD activity

The activities of QsuB and N-QsuB were determined by the change in absorption at 290 nm and calculated using a molar extinction coefficient for 3,4-DHBA (ε) equal to 3.89*10^3^ M^−1^cm^−1^ [24]. The reactions were mixed in 1 ml volumes and monitored at room temperature for 1 min.

To define the catalytic properties, a reaction was performed with 0.1 M Tris/HCl buffer (pH 7.5), 10 mM CoCl_2_, 0.1 - 5 mM DHS, and 1.5 - 2 μg of purified protein. All measurements were reproduced at least 3 times. V_max_ and K_m_ were obtained by plotting the graph in double reciprocal coordinates.

Protein inhibition studies were performed by adding 3,4-DHBA at various concentrations to the reaction mixture including 0.1 M Tris/HCl buffer (pH 7.5), 10 mM CoCl_2_, 1 mM DHS, and 1.5 - 2 μg of purified protein.

The metal cofactors were determined by monitoring the reaction in the presence of 1 mM DHS and 10 mM of the following compounds: CaCl_2_, CoCl_2_, MgCl_2_, MnCl_2_, FeCl_2_, CuSO_4_ and ZnSO_4_. The pH of the Tris/HCl buffer was varied from 7 to 8.8 in the reaction mixture in the presence of 1 mM DHS and 10 mM CoCl_2_ to determine the optimal pH.

The dioxygenase activity of the C-terminal domain was tested with the crude extracts in a reaction mixture of 0.1 M Tris/HCl buffer (pH 7.5), 10 mM MgCl_2_, 0.1 mM DHBA, and 2 μg of overall protein.

### Sequence alignment and 3D structural analysis

The QsuB sequence was aligned with QuiC1 using T-Coffee software [25]. The X-ray crystal structure of *P. aeruginosa* DHD QuiC1 was downloaded from the PDB database (http://www.rcsb.org/pdb/home/home.do). The 3D structure of QsuB was predicted using I-TASSER software [26].

## Results and discussion

### Recombinant overexpression and oligomeric state determination of QsuB and its DSD domain

According to the protein sequence alignment, QsuB had 39.9% identity with QuiC1 and 50.9% identity in the N-terminal domain coding for DSD activity. This level of identity allowed us to predict the 3D structure of QsuB based on the QuiC1 crystal structure (PDB ID: 5HMQ) (Fig 2). All the residues (Arg64, Leu97, Glu134, Leu136, His168, Asp165, Gln191, Ser206, Arg210 and Glu239) identified previously in the active site of DSD of QuiC1 [14] were also present in the N-terminal domain of QsuB.

**Fig 2.**
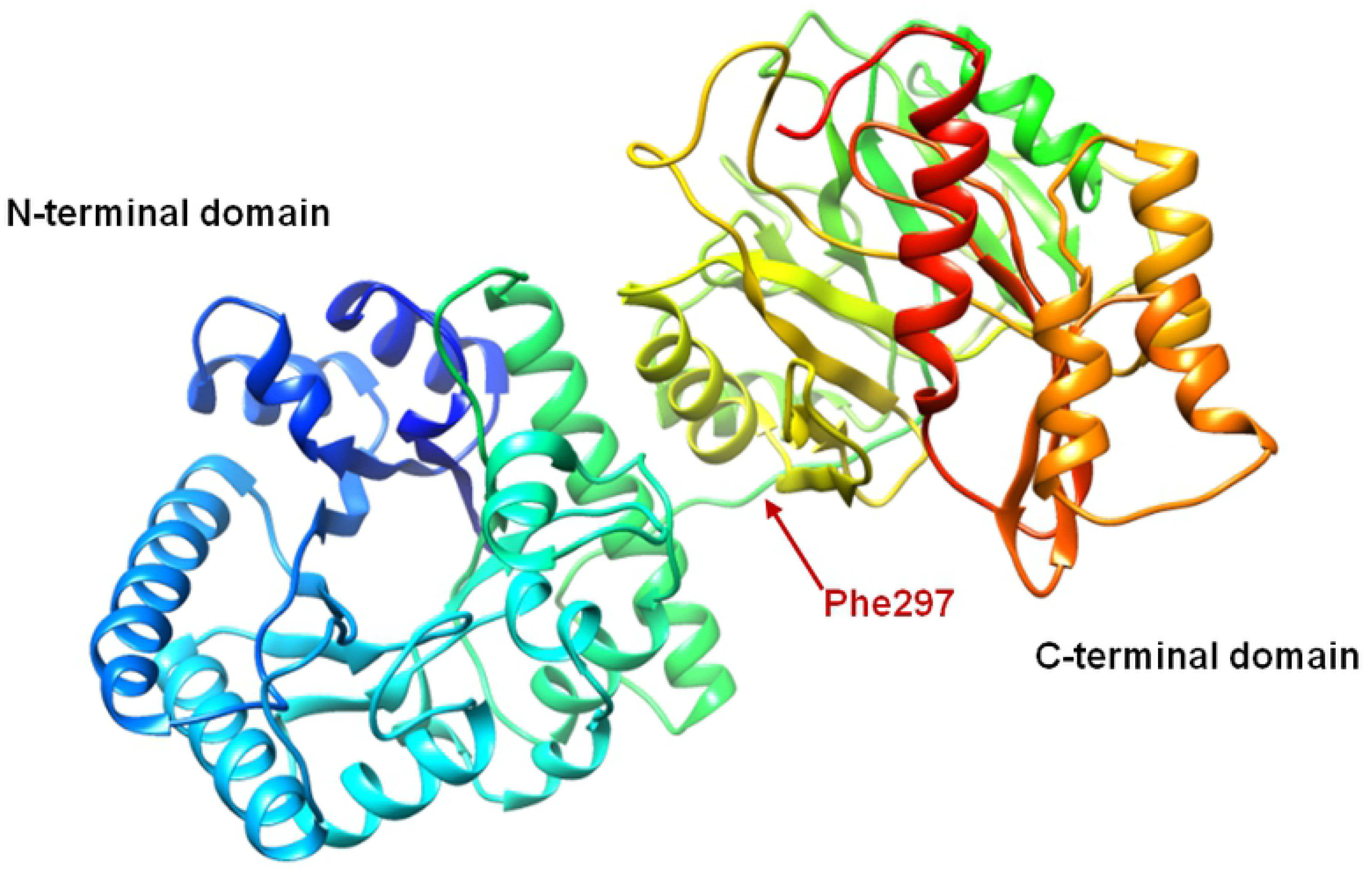
Predicted structure of the QsuB monomer. The arrow indicates where QsuB was truncated to obtain the N-QsuB variant.

To investigate the importance of both domains in 3,4-DHBA synthesis, QsuB and its N-terminal domain (N-QsuB) were overexpressed in *E. coli* BL21(DE3) cells. The full-length protein and its truncated variant were clearly visible on the electropherograms of the cell extract proteins, where they accounted for approximately 15% of the total cellular proteins (Fig 3).

**Fig 3.**
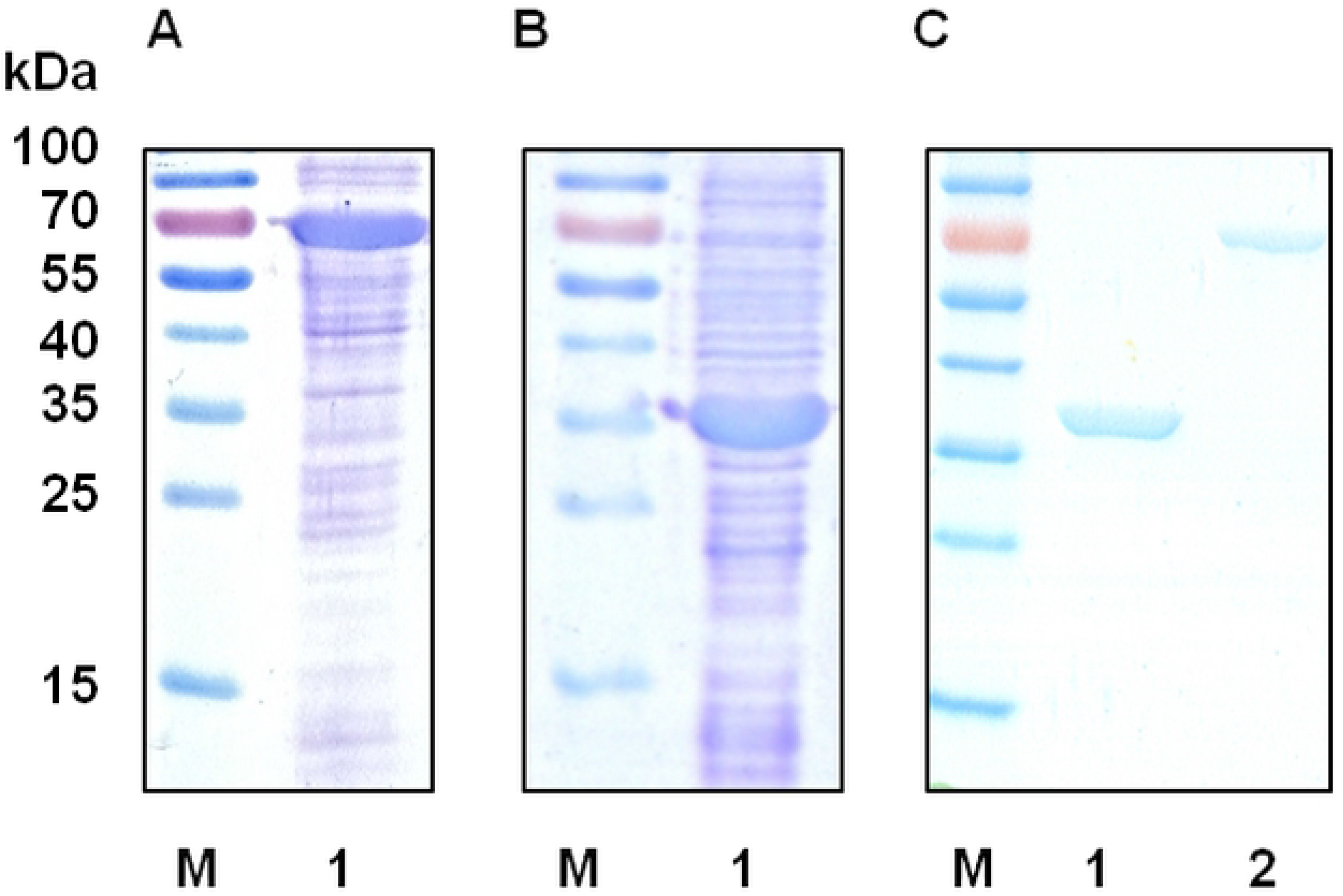
Overexpression of QsuB and N-QsuB in the T7 system. Extracts of cells after IPTG induction were analyzed using 12% SDS-PAGE. M, protein marker. (A) Lane 1, crude extract of BL21(DE3)/pET22b-*qsuB* cells. (B) Lane 1, crude extract of BL21(DE3)/pET22b-*n*-*qsuB* cells. (C) Proteins after purification: lane 1, N-QsuB; lane 2, QsuB.

**Fig 4.**
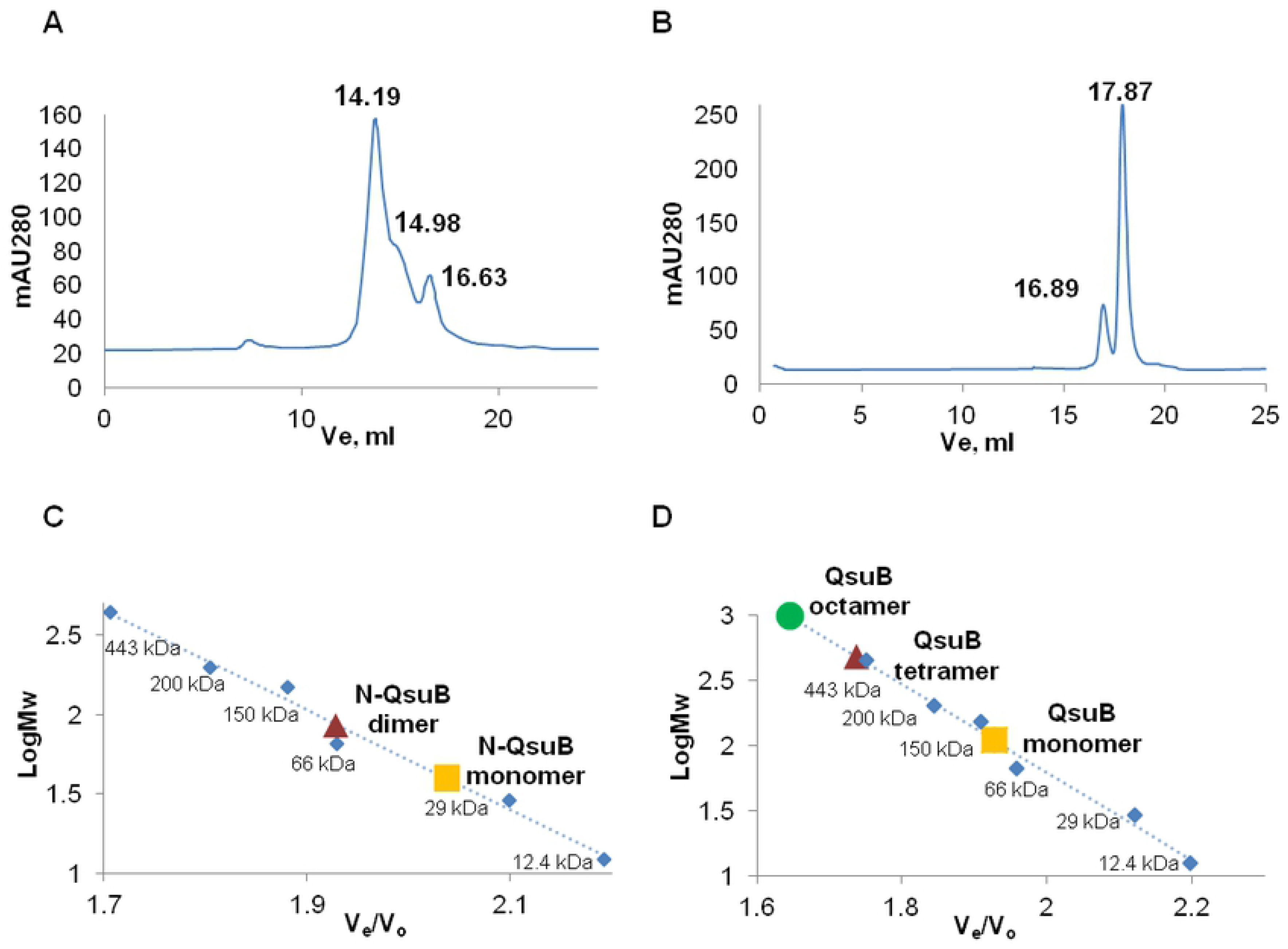
Oligomeric state determination of QsuB and N-QsuB. Molecular mass determination was carried out using SEC. Proteins were detected by monitoring absorbance at 280 nm. (A) Gel-filtration analysis of QsuB: the volumes of elution (V_e_) for the monomeric, tetrameric and octameric forms were 16.63 ml, 14.98 ml and 14.19 ml, respectively. (B) Gel-filtration analysis of N-QsuB: the V_e_ values of the monomeric and dimeric forms were 16.89 ml and 17.87 ml, respectively. The molecular mass calibration curves for QsuB (C) and N-QsuB (D) represent A_280 nm_, which is dependent on V_e_/V_o_, where V_0_ was the void volume of the column. V_0_ was determined experimentally as the V_e_ of Blue Dextran (2000 kDa). The standard proteins are represented with blue rhombuses and are cytochrome с (12.4 kDa), carbonic anhydrase (29 kDa), albumin (66 kDa), alcohol dehydrogenase (150 kDa), β-amylase (200 kDa), apoferritin (443 kDa) and thyroglobulin (669 kDa).

The purified proteins (Fig 3C) were used in gel-filtration experiments. Analysis by size exclusion chromatography (SEC) indicated that QsuB tended to form multimeric structures: it occurred as 60% octamers, 25% tetramers and 15% monomers in solution. N-QsuB was present mainly in monomeric form (80%). Thus, the C-terminal domain apparently stimulated the formation of oligomeric structures. Unlike QsuB, QuiC1 was a dimer in solution according to SEC. However, in each QuiC1 dimer, the N-terminal domain associated mainly with the C-terminal domain of the adjacent molecule [14]. Therefore, QsuB could form oligomers in the same manner as QuiC1: via interactions of the N- and C-terminal domains of adjacent units.

### Effects of metal ions and pH on the DSD activity of QsuB

Both the purified enzymes (QsuB and N-QsuB) possessed DSD activity, which was primarily detected in the presence of Mg^2+^. To investigate metal cofactor specificity, the enzymes were also tested for catalytic activity in the presence of Ca^2+^, Co^2+^, Cu^2+^, Fe^2+^, Mg^2+^, Mn^2+^ and Zn^2+^. As shown in Fig 5, Co^2+^ provided the maximal activity, followed by Mg^2+^, Mn^2+^ and Ca^2+^ in descending order. Both enzymes had almost undetectable activity in the presence of Fe^2+^, Zn^2+^ and Cu^2+^. The activity of N-QsuB in all cases was slightly less than that of QsuB. Notably, Co^2+^ is the main cofactor of QuiC1 [16], while Mn^2+^ is the main cofactor of AsbF [13]. Thus, QsuB was more similar to QuiC1 based on cofactor specificity.

**Fig 5.**
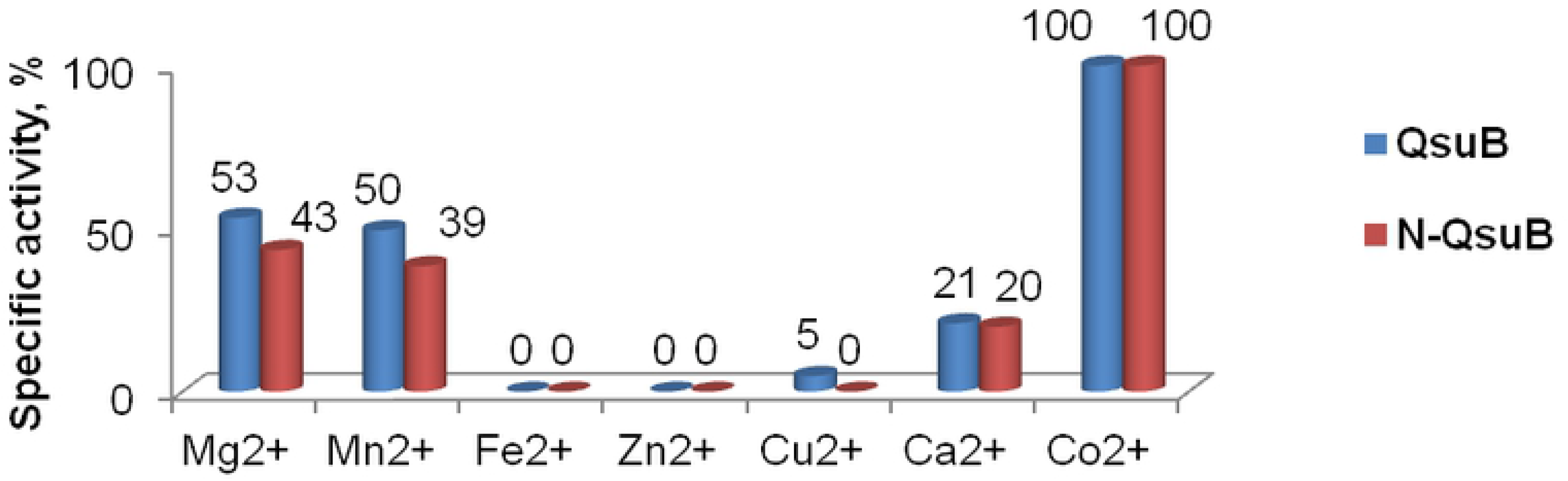
Effect of metal ions on QsuB and N-QsuB DSD activity.

The pH dependence of QsuB and N-QsuB was investigated in the presence of Co^2+^ (Fig 6). The optimum pH for these enzymes was 8.0-8.4. QsuB demonstrated a typical bell-shaped curve, while the curve of N-QsuB was smoother. Both the changed stability of the truncated protein N-QsuB and the absence of activity associated with the C-terminal domain could influence the shape of the activity curve depending on pH.

**Fig 6.**
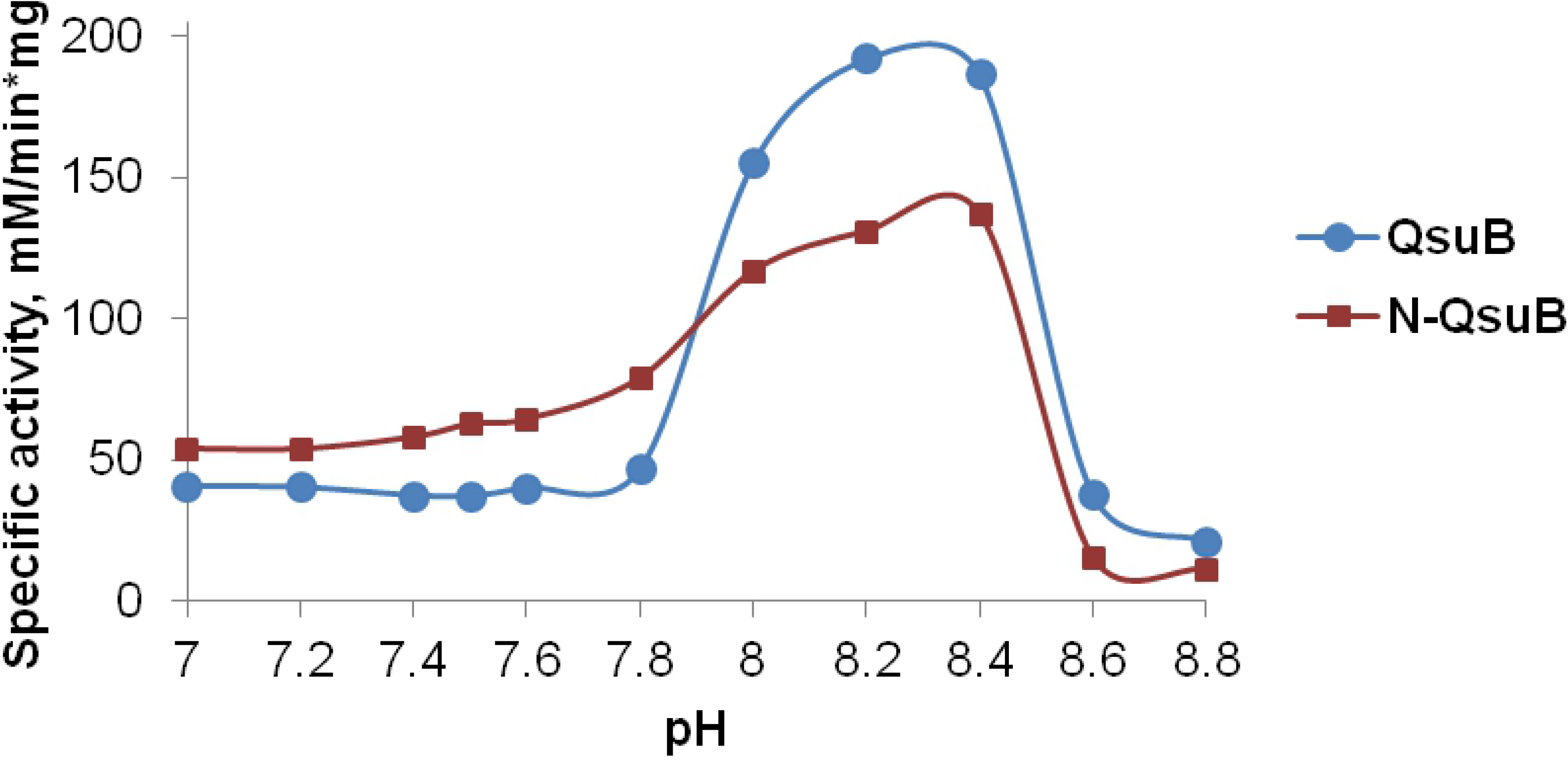
Effect of pH on QsuB and N-QsuB DSD activity.

In particular, the dioxygenase activity could manifest at those points in the curves of Fig 6 where the activity of N-QsuB, measured as the amount of 3,4-DHBA formed, was higher than the activity of QsuB. The activity of N-QsuB was higher than that of QsuB in the pH range from 7 - 7.9. This could mean that some amount of 3,4-DHBA was transformed into β-CMA with the help of QsuB under these conditions due to dioxygenase activity of the C-terminal domain. The formation of β-CMA could be detected by UV spectroscopy [27]. The QsuB protein was incubated at pH 7.5 in the presence of 3,4-DHBA for up to 60 min, but no difference in the UV spectrum was observed (Fig 7). Therefore, 3,4-DHBA dioxygenase activity was not detected under our experimental conditions.

**Fig 7.**
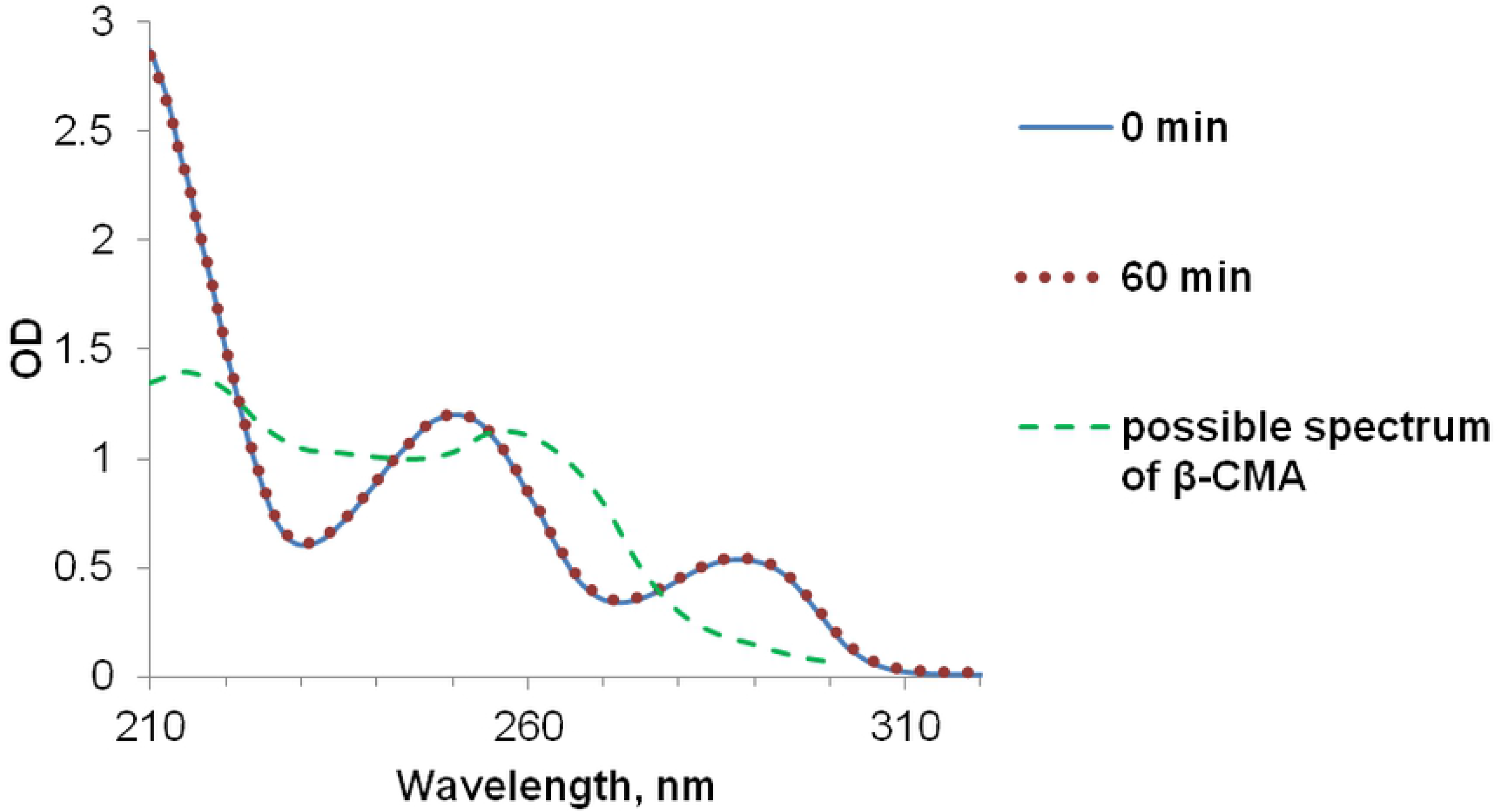
Stability of 3,4-DHBA in the presence of QsuB. The UV spectrum of the reaction mixture (pH 7.5) containing purified QsuB before and after 60 min incubation with 3,4-DHBA was the same and coincided with the 3,4-DHBA spectrum. The UV spectrum of β-CMA is shown in [27].

### Kinetic properties of QsuB and N-QsuB

Kinetic curves were obtained for QsuB and N-QsuB (Fig 8) at physiological pH for *C. glutamicum* and *E. coli*, which is 7.5 [28]. The curves seemed to demonstrate Michaelis-Menten kinetics up to 3 mM and 4 mM DHS for QsuB and N-QsuB, respectively. For this range, the K_*m*_ and k_*cat*_ parameters were calculated (Table 1). Under higher DHS concentrations, QsuB and N-QsuB activities decreased. This meant that they were inhibited by the substrate or the product formed. As seen from Fig 8 and Table 1, the removal of the C-terminal domain affected the catalytic properties of QsuB, as N-QsuB had lower K_m_ and k_cat_ values. The affinity of QsuB for DHS was lower than that of QuiC1 and much lower than that of AsbF. However, QsuB had a significantly higher turnover rate (k_*cat*_ ~ 61 s^−1^) than AsbF.

**Table 1.** Kinetic properties of QsuB and N-QsuB.

| Protein | $K_m(\text{DHS}), \mu\text{M}$ | $k_{cat}, \text{s}^{-1}$ | Source |
| --- | --- | --- | --- |
| QsuB | $961 \pm 77$ | $60.8 \pm 0.9$ | This work |
| N-QsuB | $473 \pm 71$ | $23.4 \pm 0.4$ | This work |
| QuiC1 | $331 \pm 51$ | $163.6 \pm 8.5$ | [16] |
| AsbF | $5 \pm 1$ | $0.5 \pm 0.1$ | [12] |

**Fig 8.**
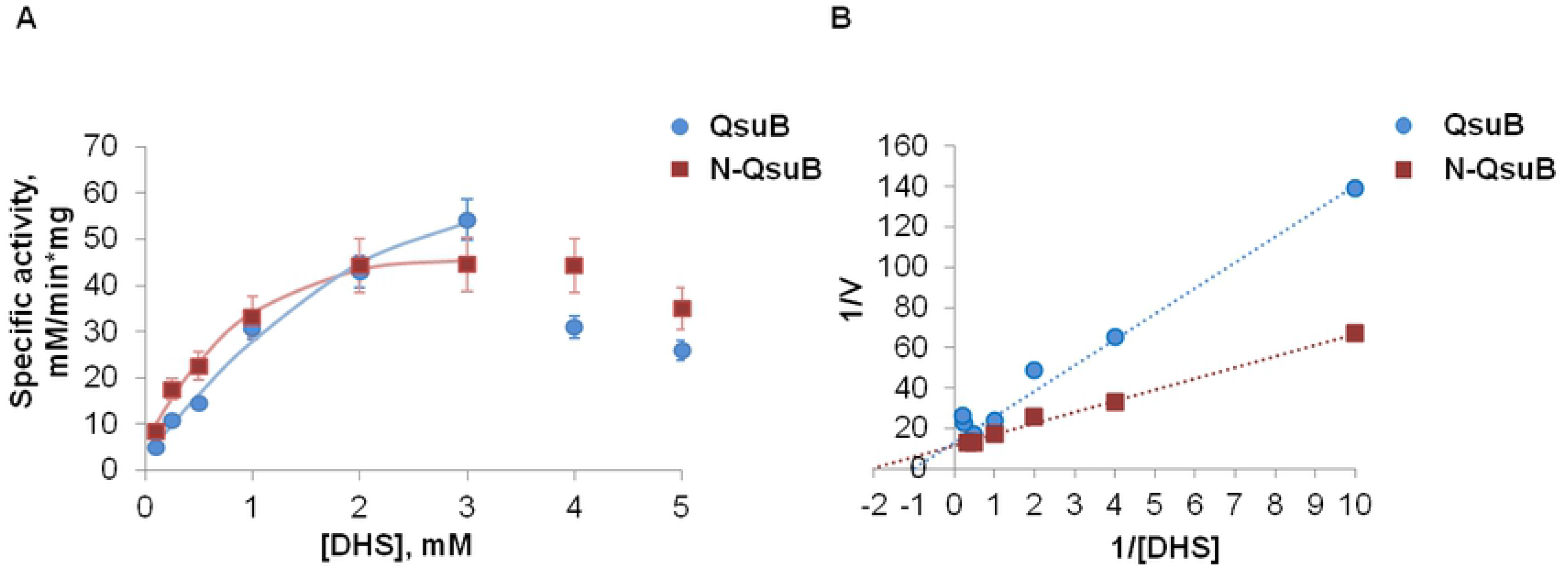
Kinetic curves of QsuB and N-QsuB. (A) Dependence of protein activity on substrate concentration. (B) Double reciprocal plots of protein kinetics.

To investigate inhibition by the product, Lineweaver–Burk plots for QsuB and N-QsuB were built (Fig 9) and showed that 3,4-DHBA tended to inhibit both enzymes. The double reciprocal coordinate graphs were linear and parallel, indicating uncompetitive inhibition. Thus, 3,4-DHBA bound to the enzyme-substrate complex (ES) and did not bind to the enzyme. As seen from the curves, 3,4-DHBA decreased both the K_*m*_ for DHS and the V_*max*_. The inhibitor binds to and stabilizes the ES, making it more difficult for S to dissociate or to be converted to product, increasing enzyme affinity for S and reducing substrate K_*m*_. Under the highest DHS concentrations, near the axis line, the enzyme was probably inhibited by the substrate as well.

**Fig 9.**
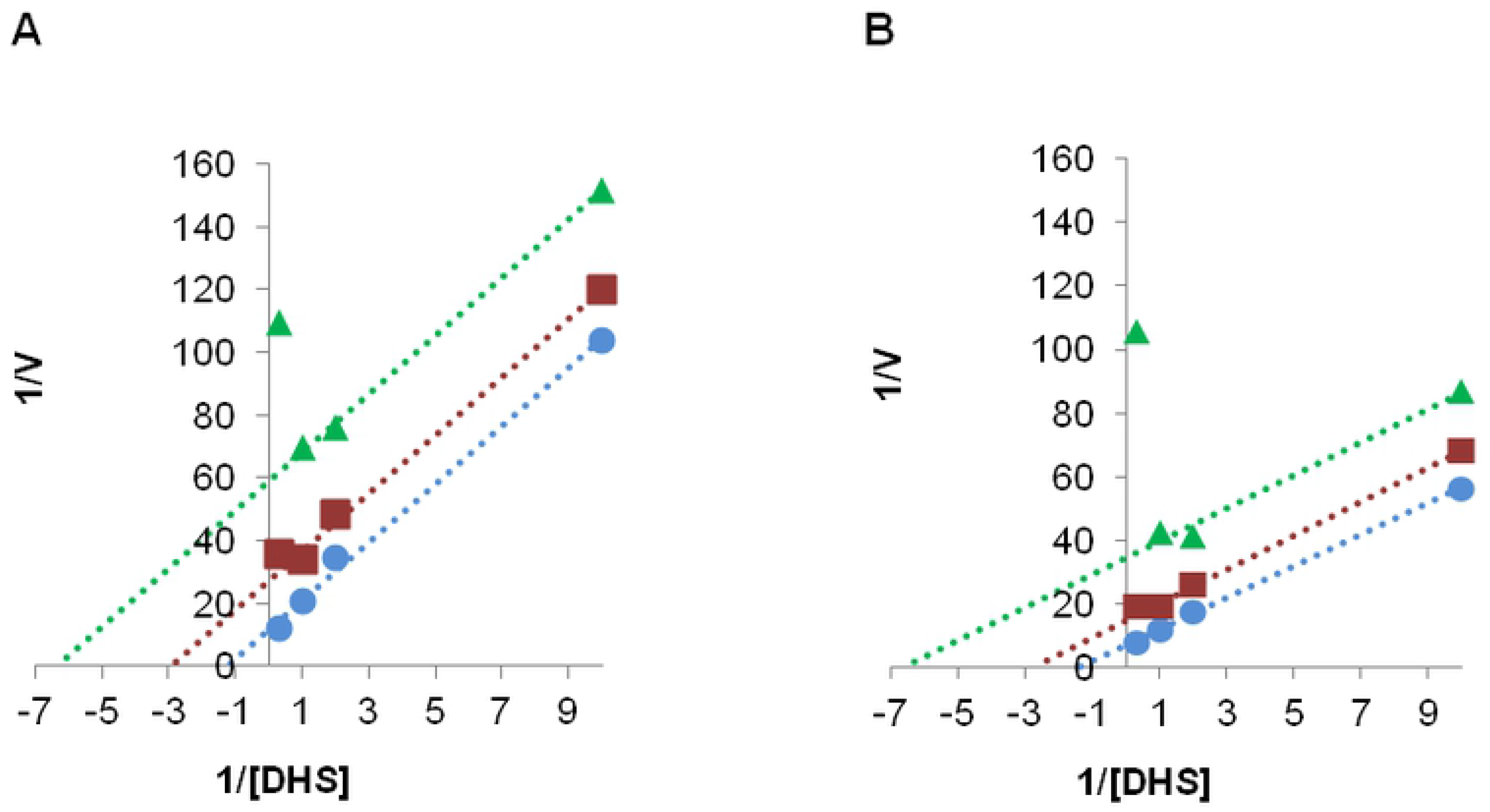
Double reciprocal plots demonstrating QsuB (A) and N-QsuB (B) inhibition. Different 3,4-DHBA concentrations are represented by colors: 0 (blue line), 0.3 (red line), and 0.6 mM (green line).

### Production of 3,4-DHBA from glucose via QsuB DSD activity

*E. coli* does not degrade 3,4-DHBA, so the DSD activity of QsuB was tested *in vivo* in *E. coli* cells. The genes *qsuB* and *N-qsuB* under the control of the IPTG-inducible promoter P_*lac*_ were integrated into the chromosome of *E. coli* MG1655Δ*aroE*. Inactivation of the shikimate dehydrogenase encoded by the *aroE* gene provided DHS accumulation in the parent strain. The obtained strains were tested for 3,4-DHBA production by fermentation with glucose as the carbon source (Table 2).

**Table 2.** Production of DHS and 3,4-DHBA from glucose (40 g/l) in test tubes.

| MG1655$\Delta aroE$<br>strain | OD<sub>540</sub> | DHS, g/l | 3,4-DHBA,<br>g/l | Res. glucose,<br>g/l | IPTG,<br>1 mM |
| --- | --- | --- | --- | --- | --- |
| — | 25±1 | 2.7±0.2 | < 0.1 | 7±1 | — |
|  | 25±3 | 2.4±0.2 | < 0.1 | 6±1 | + |
| $P_{lac}$-*qsuB* | 27±3 | 2.0±0.2 | 0.9±0.2 | 7±1 | — |
|  | 27±1 | 0.1±0.1 | 2.2±0.1 | 8.5±0.2 | + |
| $P_{lac}$-*n-qsuB* | 27±1 | 2.5±0.1 | < 0.1 | 8±1 | — |
|  | 26±2 | 1.1±0.1 | 1.3±0.1 | 7.3±0.2 | + |

As seen from Table 2, the MG1655Δ*aroE* strain began to produce 3,4-DHBA after the introduction of the P_*lac*_-*qsuB* or P_*lac*_-*n-qsuB* gene. The accumulation of 3,4-DHBA was higher in the MG1655*ΔaroE* P_lac_-*qsuB* strain. This strain produced 3,4-DHBA with and without IPTG addition, although the production was two times higher in the latter case. The MG1655*ΔaroE* P_*lac*_-*n-qsuB* strain produced 3,4-DHBA only in the presence of IPTG, and its accumulation was nearly two times less than that in the MG1655*ΔaroE* P_*lac*_-*qsuB* strain under the same conditions. Thus, the decrease in the activity of the truncated protein N-QsuB was reproduced in the *in vivo* test.

## Conclusions

Our investigations showed that QsuB was present mainly in octameric form in solution, and this form seemed to be the most active. N-QsuB had reduced activity *in vitro* and *in vivo*. The major fraction of N-QsuB in solution occurred in a monomeric form, and only a minor fraction had a dimeric form. Therefore, the C-terminal domain could be involved in oligomerization. Indeed, QuiC1, a homolog of QsuB, also has a multimeric structure that includes six protomers in an asymmetrical unit [14].

Despite the production of 3,4-DHBA from glucose using DSDs of different types, the inhibition characteristics of such enzymes by 3,4-DHBA were not investigated. The accumulation of 3,4-DHBA using QsuB was demonstrated in *E. coli* cells containing only one modification, Δ*aroE*. In addition, 3,4-DHBA production was reversibly correlated with DHS as a byproduct. Even under the strongest *qsuB* gene induction by IPTG, residual DHS was found. This byproduction of DHS could be due to both enzyme deficiency and feedback inhibition. QsuB was inhibited by the product 3,4-DHBA via uncompetitive inhibition. This result indicates that 3,4-DHBA could affect only the ES and not the free enzyme. Biosynthetic enzymes with this type of inhibition are rare in nature and are thought to have been lost during evolution [29]. Usually, uncompetitive inhibition is irreversible, and it provides very attractive prospects for drug design [29,30].

Most likely, QsuB is more suitable for obtaining 3,4-DHBA as an intermediate product. Studying the inhibition of other DSDs by 3,4-DHBA as well as obtaining suitable mutant forms of QsuB are two alternative ways to select an appropriate enzyme for large-scale 3,4-DHBA production.

## Supporting information

**S1 Table. Primers used in investigation.**

## Acknowledgments

We are grateful to Dr. Nataliya V. Stoynova for critical reading the manuscript.

## Author Contributions

**Conceptualization:** VGD VOV EAS.

**Formal analysis:** VOV EAS.

**Investigation:** EAS VOV MVO SVS.

**Methodology:** VOV EAS VGD.

**Supervision:** VGD.

**Validation:** EAS MVO.

**Visualization:** EAS SVS.

**Writing ± original draft:** EAS.

**Writing ± review & editing:** VGD EAS.

